# Transcriptomics and Metabolomics Reveal Tomato Consumption Alters Hepatic Xenobiotic Metabolism and Induces Steroidal Alkaloid Metabolite Accumulation in Mice

**DOI:** 10.1101/2023.04.18.536606

**Authors:** Michael P. Dzakovich, Mallory L. Goggans, Jennifer M. Thomas-Ahner, Nancy E. Moran, Steven K. Clinton, David M. Francis, Jessica L. Cooperstone

## Abstract

**Scope:** Tomato consumption is associated with many health benefits including lowered risk for developing certain cancers. It is hypothesized that tomato phytochemicals are transported to the liver and other tissues where they alter gene expression in ways that lead to favorable health outcomes. However, the effects of tomato consumption on mammalian liver gene expression and chemical profile are not well defined.

**Methods and results:** We hypothesized that tomato consumption would alter mouse liver transcriptomes and metabolomes compared to a control diet. C57BL/6 mice (n=11-12/group) were fed a macronutrient matched diet containing either 10% red tomato, 10% tangerine tomato, or no tomato powder for 6 weeks after weaning. RNA-Seq followed by gene set enrichment analyses indicated that tomato type and consumption, in general, altered expression of phase I and II xenobiotic metabolism genes. Untargeted metabolomics experiments revealed distinct clustering between control and tomato fed animals. Nineteen molecular formulas (representing 75 chemical features) were identified or tentatively identified as steroidal alkaloids and isomers of their phase I and II metabolites; many of which are reported for the first time in mammals.

**Conclusion:** These data together suggest tomato consumption may impart benefits partly through enhancing detoxification potential.

## 1 Introduction

Epidemiological studies indicate that consuming tomatoes and tomato products is associated with a reduced risk of developing many chronic diseases including prevalent cancers ^[1,2]^. The principal hypothesis is that the carotenoid lycopene is responsible for these clinical outcomes ^[1–12]^. However, some studies comparing consumption of diets with tomato vs. those containing the same amount of lycopene as a purified compound, show that benefits are greater with consumption of the whole food ^[13,14]^. These findings suggest that other phytochemicals are contributing to the bioactivity of tomato in addition to lycopene.

The liver is one of the earliest destinations for absorbed dietary small molecules. Small molecules can be further metabolized in the liver prior to distribution to plasma, tissue, and/or modified for excretion via urine or bile through phase I and II metabolism. Rodent studies have established that dietary supplementation with tomato fruit extracts and specific phytochemicals can alter the progression of nonalcoholic fatty liver disease, nonalcoholic steatohepatitis, and hepatocellular carcinoma partially through changes in gene expression ^[8–11,15]^. Prior investigations into the effects of tomato consumption on health have relied on targeted gene expression and phytochemical analyses (e.g. carotenoids) ^[16–18]^. While prior investigations have tested singular tomato varieties, it is noteworthy that tomato composition can vary drastically based on genotype and pre/post-harvest environmental conditions^[19]^. Biological responses to one tomato containing intervention may not be reflective of responses to tomato, in general. Feeding studies using multiple tomato varieties, next generation sequencing, and untargeted metabolomics offer a more comprehensive alternative approach to examine and discover relationships between tomato consumption and health outcomes. Utilizing these approaches in concert may reveal novel phytochemicals of interest and indicate potential mechanisms by which tomato consumption affects health.

This study aimed to define the impact of tomato consumption on mouse liver transcriptional and chemical profiles using RNA-Seq and untargeted mass spectrometry-based metabolomics. Mice were fed diets containing one of two distinct tomato varieties to resolve responses to tomato consumption, in general, from those due to specific tomato phytochemicals unique to a given cultivar, or a macronutrient matched, tomato-free control diet. Untargeted metabolomic analyses were focused on the more chemically diverse polar/semi-polar compounds since previous studies have investigated differences due to more non-polar carotenoid profiles^[16–18]^.

## 2 Experimental Section

### 2.1 Reagents and standards

LC-MS Optima grade acetonitrile (0.1% formic acid), isopropanol, methanol, and water (0.1% formic acid), and 99.5% butylated hydroxytoluene (BHT), were purchased from Fisher Scientific (Pittsburgh, PA. USA). Alpha-tomatine was purchased from Extrasynthese (Genay, France) and tomatidine from Sigma Aldrich (St. Louis, MO, USA).

### 2.2 Animal Diets and Experimental Design

The tangerine processing tomato variety FG04-167 (enriched in *cis*-structured lycopene and its precursors due to the *t^3183^* allele ^[20,21]^) and the red processing varieties OH8243, OH8245, and FG99-218 (elevated all-*trans*-lycopene concentration due to the *old gold crimson* (*og^c^*) allele ^[22]^) were grown, processed into single strength juice, and freeze-dried as previously described ^[16,21]^. Animal diets were manufactured by Dyets (Bethlehem, PA, USA) by combining tomato powders into a modified AIN-93G diet at 10% (*w*/*w*) prior to pelleting ^[23]^. To maintain macronutrient composition across all treatments, control diets were supplemented with corn starch and dextrose to compensate for the carbohydrates naturally present in tomato ^[13]^. Animal diets were stored at −20 °C and mice received fresh diet every 2-3 days to minimize phytochemical degradation.

### 2.3 Experimental Design

At four weeks of age, male C57BL/6 mice were randomly assigned treatment groups (n=11-12/group) and fed either a control diet, a diet supplemented with 10% (*w*/*w*) red tomato powder, or a diet supplemented with 10% (*w*/*w*) tangerine tomato powder (protocol #2010A00000095). Mouse bodyweights and food intake were determined weekly to ensure consistent growth and development throughout the experiment. Final body and liver masses are reported in Table 1 with additional metadata in Supporting Table 1. Mice were sacrificed after 6 weeks of dietary intervention and one liver lobe from each mouse was split for chemical profiling or transcriptomic analysis (preserved in RNA*later* (Thermo Fisher Scientific; Waltham, MA, USA)). Samples were flash frozen in liquid nitrogen and stored at −80 °C until analysis.

**Table 1.** Mouse body and liver mass (mean ± standard deviation) at study termination (10 wk).

| Treatment<sup>a</sup> | Body Mass<br>(g) | Liver Mass<br>(g) |
| --- | --- | --- |
| Control | 28.13 ± 3.54 | 1.09 ± 0.16 |
| Red | 27.21 ± 4.08 | 1.20 ± 0.22 |
| Tangerine | 25.91 ± 3.27 | 1.15 ± 0.19 |

<sup>a</sup>ANOVA among treatment determined that the effect of diet type was non-significant for body mass ( $P = 0.35$ ) and liver mass ( $P = 0.43$ ).

### 2.4 RNA Extraction

Mouse liver RNA was extracted from 10 mg (± 2 mg) of preserved tissue using RNEasy kits and QIAshredders (QIAGEN; Germantown, MD, USA). Aliquots of RNA from each sample were assessed on an Agilent 2100 Bioanalyzer (Agilent Technologies; Santa Clara, CA, USA) and RNA Integrity Number scores ranged between 8.7 and 9.3.

### 2.5 cDNA Library Preparation and RNA-Seq Data Acquisition

Libraries were prepared using the NEXTflex Rapid Directional RNA-Seq kit (PerkinElmer; Waltham, MA, USA) and sequenced by The Cleveland Clinic Lerner Genomics Core. Libraries were sequenced on two Illumina HiSeq2500 lanes (Illumina; San Diego, CA, USA) using 100 bp paired-end reads at a depth of ∼30 million reads/sample and concurrently run with the PhiX quality control library (Illumina).

### 2.6 Analysis of RNA-Seq Data

FASTQ files were checked for quality using FASTQC through the Ohio Supercomputer Center ^[24]^. Adapter sequences and low quality reads were trimmed using Trimmomatic ^[25]^. Trimming parameters can be found in Supporting Table 2. Reads were aligned to an index of the mouse genome (GRCm38.p6) using Rsubread and BAMtools ^[26,27]^. Genes with greater than 0.37 counts per million (10/average library size) in at least 11 samples were retained (14,075 total; Supporting Table 3) and log2 transformed for visualization. Data were normalized to trimmed mean of M-values (TMM) and analyzed for differential expression using edgeR ^[28,29]^. Genewise negative binomial generalized linear models were used to compare treatment groups at a Benjamini-Hochberg false discovery rate corrected significance threshold of *P* ≤ 0.1 (Supporting Tables 4-7) ^[30–32]^. Gene set overlaps of differentially expressed genes from each pairwise treatment contrast (Supporting Tables 8-10) were determined with the Molecular Signatures Database (MSigDB; https://www.gsea-msigdb.org/gsea/msigdb/annotate.jsp; Hallmark and KEGG gene sets).

**Table 2.** Gene set overlaps enriched in differentially expressed genes from mice fed tomato supplemented diets.

|  | Tangerine<br>vs. Control | Red vs.<br>Control | Tangerine<br>vs. Red |
| --- | --- | --- | --- |
| Number of Genes in Comparison | 3<sup>a</sup> | 279 | 419 |
| Gene Set Name |  |  |  |
| KEGG linoleic acid metabolism | 2 | —<sup>b</sup> | — |
| KEGG retinol metabolism | 2 | — | — |
| KEGG metabolism of xenobiotics by cytochrome P450 | 2 | — | — |
| KEGG drug metabolism by cytochrome P450 | 2 | — | — |
| Hallmark xenobiotic metabolism | — | 10 | 17 |
| Hallmark interferon gamma response | — | 7 | 11 |
| KEGG arginine and proline metabolism | — | 4 | 6 |
| Hallmark IL2 STAT5 signaling | — | 9 | — |
| Hallmark G2M checkpoint | — | 8 | — |
| Hallmark oxidative phosphorylation | — | 8 | — |
| KEGG ABC transporters | — | 4 | — |
| Hallmark heme metabolism | — | 7 | — |
| Hallmark UV response up | — | 6 | — |
| Hallmark UV response down | — | 6 | — |
| KEGG circadian rhythm mammal | — | — | 6 |
| KEGG alanine aspartate and glutamate metabolism | — | — | 6 |
| Hallmark cholesterol homeostasis | — | — | 7 |
| Hallmark glycolysis | — | — | 10 |
| Hallmark KRAS signaling down | — | — | 10 |
| KEGG selenoamino acid metabolism | — | — | 4 |
| KEGG peroxisome | — | — | 6 |

<sup>a</sup>Number of differentially expressed genes present in each gene set
<sup>b</sup>Gene set overlap not detected

**Table 3.**
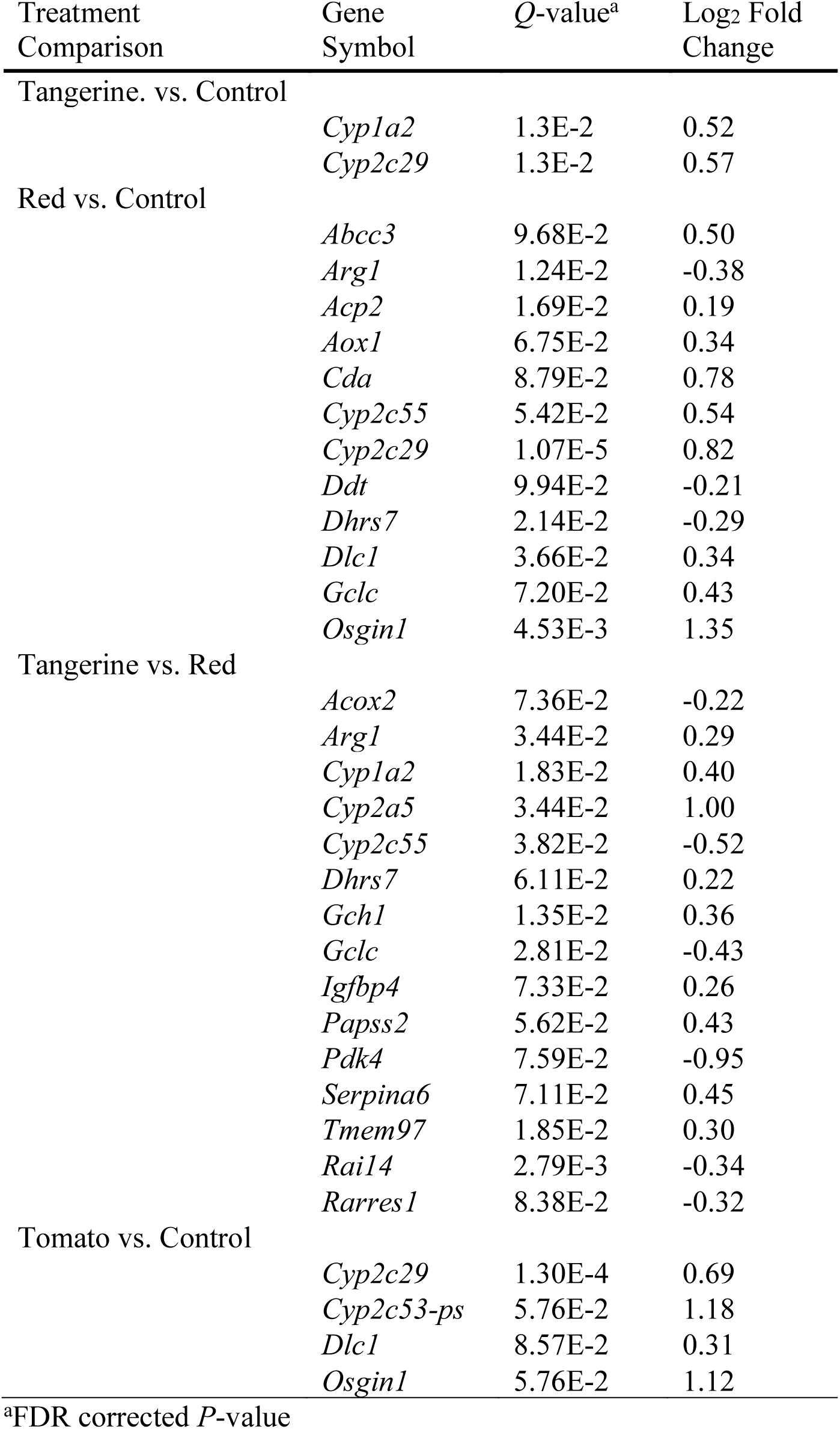
Select differentially expressed genes related to xenobiotic metabolism or disease development

| Treatment Comparison | Gene Symbol | Q-value<sup>a</sup> | Log<sub>2</sub> Fold Change |
| --- | --- | --- | --- |
| Tangerine. vs. Control |  |  |  |
|  | *Cyp1a2* | 1.3E-2 | 0.52 |
|  | *Cyp2c29* | 1.3E-2 | 0.57 |
| Red vs. Control |  |  |  |
|  | *Abcc3* | 9.68E-2 | 0.50 |
|  | *Arg1* | 1.24E-2 | -0.38 |
|  | *Acp2* | 1.69E-2 | 0.19 |
|  | *Aox1* | 6.75E-2 | 0.34 |
|  | *Cda* | 8.79E-2 | 0.78 |
|  | *Cyp2c55* | 5.42E-2 | 0.54 |
|  | *Cyp2c29* | 1.07E-5 | 0.82 |
|  | *Ddt* | 9.94E-2 | -0.21 |
|  | *Dhrs7* | 2.14E-2 | -0.29 |
|  | *Dlc1* | 3.66E-2 | 0.34 |
|  | *Gclc* | 7.20E-2 | 0.43 |
|  | *Osgin1* | 4.53E-3 | 1.35 |
| Tangerine vs. Red |  |  |  |
|  | *Acox2* | 7.36E-2 | -0.22 |
|  | *Arg1* | 3.44E-2 | 0.29 |
|  | *Cyp1a2* | 1.83E-2 | 0.40 |
|  | *Cyp2a5* | 3.44E-2 | 1.00 |
|  | *Cyp2c55* | 3.82E-2 | -0.52 |
|  | *Dhrs7* | 6.11E-2 | 0.22 |
|  | *Gch1* | 1.35E-2 | 0.36 |
|  | *Gclc* | 2.81E-2 | -0.43 |
|  | *Igfbp4* | 7.33E-2 | 0.26 |
|  | *Papss2* | 5.62E-2 | 0.43 |
|  | *Pdk4* | 7.59E-2 | -0.95 |
|  | *Serpina6* | 7.11E-2 | 0.45 |
|  | *Tmem97* | 1.85E-2 | 0.30 |
|  | *Rai14* | 2.79E-3 | -0.34 |
|  | *Rarres1* | 8.38E-2 | -0.32 |
| Tomato vs. Control |  |  |  |
|  | *Cyp2c29* | 1.30E-4 | 0.69 |
|  | *Cyp2c53-ps* | 5.76E-2 | 1.18 |
|  | *Dlc1* | 8.57E-2 | 0.31 |
|  | *Osgin1* | 5.76E-2 | 1.12 |

<sup>a</sup>FDR corrected *P*-value

### 2.7 Extraction of Polar Metabolites

Polar metabolites were extracted from 50 mg (± 5 mg) of frozen liver using a modified extraction method ^[33]^. Five, 2 mm zirconium oxide beads (Union Process; Akron, OH, USA) were placed in 1.5 mL tubes containing sample tissue and 500 μL of chilled methanol with 0.1% BHT was immediately added. Samples were homogenized using a SPEX Geno/Grinder 2010 for 30 seconds at 1400 RPM within a chilled Cryo−Block (SPEX Sample Prep; Metuchen, NJ, USA). Samples were centrifuged at 21,130 x *g* for five min at 4 °C, and 475 μL of supernatant fluid from each sample was passed through a solid-phase extraction cartridge (Oasis PRiME HLB 1cc/30mg, Waters; Milford, MA, USA) to remove phospholipids. An additional 100 μL of acetonitrile was passed through the cartridges to elute any remaining polar metabolites. Three separate 135 μL aliquots were reserved for analysis and a final 75 μL aliquot from each sample was combined to create a pooled quality control (QC) sample. Process blanks used to identify features derived from laboratory consumables were created using the same extraction, with water substituted for liver. Samples, process blanks, and QCs were dried down under a stream of nitrogen and stored at −80 °C until analysis.

### 2.8 Untargeted Metabolomics Data Collection

Extracts were redissolved in 100 μL of 1:1 methanol:water and centrifuged at 21,130 x *g* for 3 min to precipitate undissolved solids. The supernatant was run on an ultra−high performance liquid chromatography (Agilent 1290 Infinity II, UHPLC) system coupled with an quadrupole time of flight mass spectrometer (Agilent 6545, QTOF-MS). Analytes were separated on a C18 bridged ethylene hybrid column (Waters, 2.1 × 100 mm, 1.7 µm particle size), maintained at 40 °C, using water (A) and acetonitrile (B), both with 0.1% formic acid at 0.5 mL/min. The gradient used was as follows: 0% B held for 1 min, a linear gradient to 100% B over 10 min, 100% B held for 2 min, and a return to 0% B for an additional 2 min. Between injections, the needle and sample loop were washed with acetonitrile for 8 sec, isopropanol for 10 sec, and water for 5 sec to reduce carryover. Samples were maintained at 4 °C and the injection volume was 5 μL. The QTOF-MS was operated in electrospray positive ionization mode and data were collected from 50 – 1700 *m*/*z*. Gas temperature was 350 °C with a flow of 10 L/min. Nebulizer gas was 35 psig. Sheath gas temperature and flow were 375 °C and 11 L/min, respectively. To condition the instrument, a QC was repeatedly injected until base peak chromatograms stabilized. QC samples were run every 6 injections to monitor intra-run variability.

### 2.9 Analysis of Untargeted Metabolomics Data

Feature extraction and alignment were conducted in Agilent MassHunter Profinder 10.0 (Agilent Technologies). Parameters can be found in Supporting Table 11. Features were removed if they were not present in all 7 QC samples, present in <8 individuals within a treatment group, had >30% variance in QC samples, and/or had a maximum ion intensity/average intensity less than 10 times that of the process blank (Supporting Table 12). Missing values were imputed with half of the lowest peak area value in the dataset. Data were log_2_ transformed and analyzed in R 3.6.3 ^[34]^. Principal components analysis (PCA) was used for visualization using the packages FactoMineR, Factoextra, and ggplot2 ^[35,36]^. K-means clustering was conducted using R 3.6.3 (kmeans, nstart = 20, iter.max = 200). P-values from univariate statistics such as t-tests (t_test) and analysis of variance (ANOVA; anova_test) were corrected for multiple testing using the Benjamini-Hochberg procedure (adjust_pvalue) using the package rstatix ^[37,38]^.

### 2.10 UHPLC-QTOF-MS/MS Feature Identification

Fragmentation data of features of interest were collected using an Agilent 1290 Infinity II UHPLC in tandem with an Agilent 6545 QTOF-MS using pooled extracts of red and tangerine fed mouse livers (n=4/group). MS1 collection with the addition of independent MS/MS experiments were conducted as described previously with 10-20 µL injections. A collision energy of 30 eV was used for all masses and identified masses are detailed in Supporting Table 14.

### 2.11 Dataset Availability

Raw datafiles from RNA-Seq and untargeted metabolomics are publicly available in the Gene Expression Omnibus (https://www.ncbi.nlm.nih.gov/geo/query/acc.cgi?acc=GSE221230) and MetaboLights (www.ebi.ac.uk/metabolights/MTBLS6715) ^[39]^, respectively. Gene counts, differentially expressed genes, gene set overlaps, and deconvoluted metabolomics data are available in the supporting information. All code for RNAseq and metabolomics analyses can be found at www.github.com/CooperstoneLab/tomato-mouse-omics.

## 3 Results and Discussion

### 3.1 Animal weights and tissue mass

Body and liver masses of mice in each treatment group did not significantly differ following six weeks of *ad libitum* feeding (Table 1) in alignment with previous rodent and pig studies ^[16,40–42]^. Given that diets were macronutrient matched, differences seen in RNA-Seq and untargeted metabolomics datasets can be attributed to tomato consumption.

### 3.2 Input data for RNA-Seq were suitable for analysis

RNA-Seq was utilized to gain a broad perspective of the influence of tomato consumption on gene transcription in mouse liver. Figures S1A-D demonstrate that input data were suitable for analysis after normalization, did not exhibit lane effects, and dispersion was not skewed ^[43]^. In parallel with other dietary intervention studies utilizing RNA-Seq, an FDR cutoff of 0.1 was set to decrease the probability of excluding differentially expressed genes due to type II error ^[30–32]^. A total of 355 and 450 unique genes were up and downregulated, respectively, in response to diet.

### 3.3 Tomato consumption and type influences the expression of xenobiotic metabolism genes

Gene set enrichment analysis (GSEA) using MSigDB contextualized metabolic alterations modified by diet. Gene sets varied by treatment contrast, but differential regulation of xenobiotic metabolism genes was a recurrent trend (Table 2 and 3). When tangerine tomato supplemented mice were compared to mice receiving control diets, linoleic acid metabolism, retinoid metabolism, and xenobiotic and drug metabolism by cytochrome P450 enzymes gene sets were significantly enriched (Table 2). Significant upregulation of the cytochrome P450s, cytochrome P450, family 2, subfamily c, polypeptide 29 (*Cyp2c29)* and *Cyp1a2* drove this outcome (Table 3). These phase I enzymes are downregulated by high-fat diets and environmental pollutants ^[44]^ and suppressed in fatty liver disease ^[45]^. More recently, *Cyp2c29* was found to be downregulated in cancerous liver tissue and overexpression protected mice from inflammatory damage ^[46]^. Supplementation with other vegetables like broccoli has also been shown to increase expression of phase I and II metabolizing enzymes ^[47]^. It is possible that one way in which tomato consumption might be exerting positive health outcomes is by upregulating detoxifying enzymes, though this hypothesis requires further testing.

Xenobiotic metabolism genes were most frequently altered when comparing mice fed red tomato enriched diets vs. control animals (Table 2). Glutamate-cysteine ligase (*Gclc*), for example, encodes a protein that generates the precursor for glutathione; a molecule bonded to xenobiotics to aid in excretion ^[48]^. Other genes such as dehydrogenase/reductase 7 (*Dhrs7*) were decreased in response to tomato consumption, however the implications of this finding are unclear.

Comparing mice fed diets supplemented with red or tangerine tomatoes revealed multiple low-magnitude increases in xenobiotic metabolism genes including multiple cytochrome P450s (*Cyp1a2, Cyp2a5,* and *Cyp2c55*) as well as 3′-phosphoadenosine 5′-phosphosulfate synthase 2 (*Papss2*; Table 3). These P450s participate in phase I xenobiotic and endogenous steroid metabolism while *Papss2* encodes a phase II sulfonation enzyme ^[49,50]^. By contrast, other xenobiotic metabolism genes were downregulated including retinoid acid induced 14 (*Rai14*) and retinoic acid receptor responder 1 (*Rarres1*). We hypothesize their downregulation may be due to feedback inhibition in vitamin A sensing caused by carotenoid metabolites unique to tangerine tomatoes ^[51]^.

We reclassified our data combining mice that consumed diets containing red or tangerine tomato into one group (“tomato”) to better understand the effect of general tomato consumption. No significant gene set overlaps were detected by GSEA in this contrast. Of the genes that were differentially expressed and upregulated (Supporting Table 7), *Cyp2c29*, oxidative stress induced growth inhibitor 1 (*Osgin1*), and deleted in liver cancer 1 (*Dlc1*) have been shown to be downregulated or knocked out entirely during the development of hepatocellular carcinoma (HCC) ^[46,52,53]^. It is unclear if the protective effects associated with tomato consumption are due in part to the upregulation of these and other genes as we observed in our study.

Aside from xenobiotic metabolism, many other contrast-specific gene sets were detected (Table 2). Several mammalian circadian rhythm genes were differentially expressed when comparing red to tangerine tomato supplemented mice (Supporting Table 6 and 10). Tomato-consuming mice with HCC or early prostate carcinogenesis are susceptible to alterations in circadian rhythm genes including period circadian regulator 2 (*Per2*), clock circadian regulator (*Clock*), cryptochrome circadian regulator 2 (*Cry2*), and aryl hydrocarbon receptor nuclear translocator-like protein 1 (*Arntl*) ^[54,55]^. However, expression differences did not manifest themselves in healthy, tomato-supplemented animals ^[55]^. Similarly, circadian rhythm gene expression in the current study was not different between tomato-fed versus control animals. The implications of differentially expressed circadian rhythm genes when comparing red and tangerine tomato supplemented mice are unclear.

Previous studies in rodents using PCR-based assays demonstrated that tomato and/or the lycopene consumption have marked effects on the development of diseases leading to and including HCC by altering gene expression signatures ^[9–11]^. While each contrast in our own study had unique signatures, genes associated with mammalian xenobiotic metabolism tended to be the only consistent category of differentially expressed genes. We conducted an untargeted metabolomics analysis to determine how tomato consumption affected hepatic xenobiotic metabolism.

### 3.4 Input data for untargeted metabolomics were suitable for analysis

After data pre-processing, 2,133 chemical features were retained for analysis. A boxplot of log_2_ transformed feature abundance by sample demonstrates consistent measurement of identical QCs and good instrumental reproducibility (Figure S2A). Principal components analysis with all samples and features, including QCs, was also conducted to assess data quality (Figure S2B). Close clustering of QCs indicates our data are not confounded by run order, and that identical samples can be reproducibly measured.

### 3.5 The chemical landscape of mouse liver tissue is altered by tomato consumption

Data were analyzed both by comparing the three treatment groups (red vs. tangerine vs. control) as well as tomato vs. control. A PCA scores plot shows distinct separation between control samples and those from mice fed diets supplemented with red or tangerine tomatoes on PC1 (Figure 1). Samples from mice consuming diets supplemented with red or tangerine tomato were not visually separated, suggesting no overwhelming differences in polar/semi-polar molecules. We also used k-means clustering as an unsupervised analysis to heuristically investigate natural clusters in our data, and how these clusters compare to what we observe using PCA. PCA scores plots colored by k-means assignment with 2 and 3 clusters are Figures S2C and S2D, respectively. In Figure S2C, k-means clustering distinguishes without error between animals fed tomato vs. control diets. Misclassification can be clearly observed in Figure S2D when attempting to distinguish 3 clusters (individual tomato varieties vs control). This finding indicates more distinguishing features differ between mice fed a control diet or those fed diets enriched with tomato. Because of more easily distinguishable differences between tomato and control, we focused our analysis between these groups.

**Figure 1.**
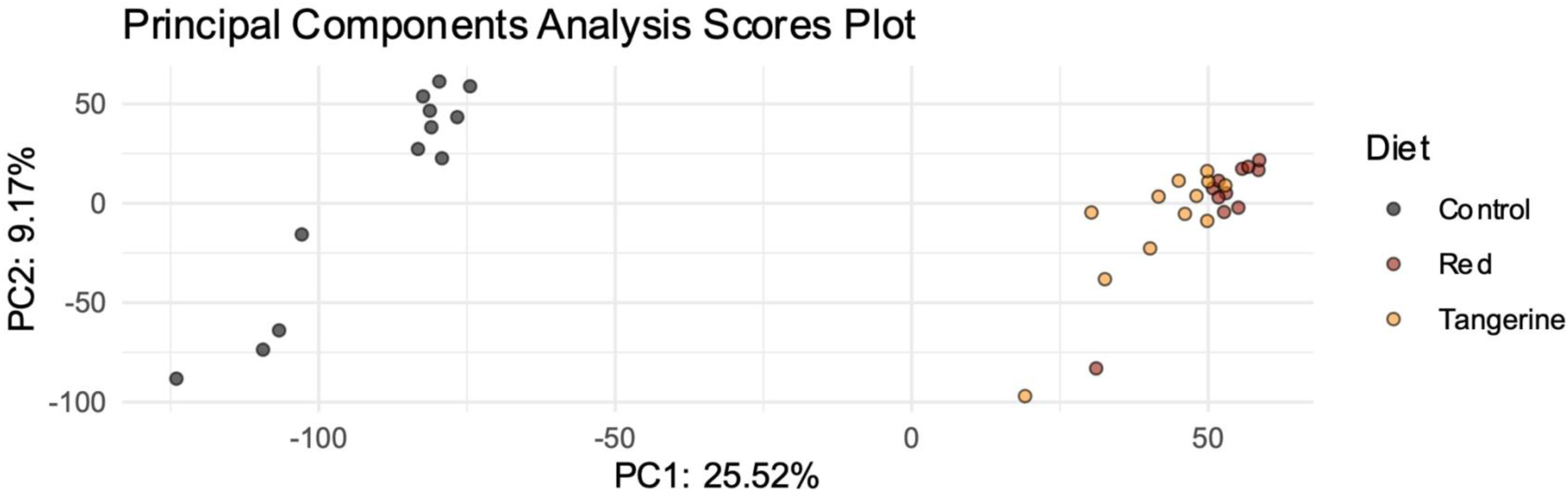
Principal components analysis scores plots visualizing data structure from untargeted metabolomics (ESI+) data excluding quality control samples.

Outcomes from univariate t-tests comparing tomato supplemented animals to controls are displayed as a volcano plot to visualize statistical significance against change in abundance (Figure 2). Most differentially present features were increased in tomato fed vs control (Figure 2, red points) indicating these features are likely tomato phytochemicals and/or their metabolites. High significance values of these features bolster the hypothesis that these compounds are derived from tomato consumption.

**Figure 2.**
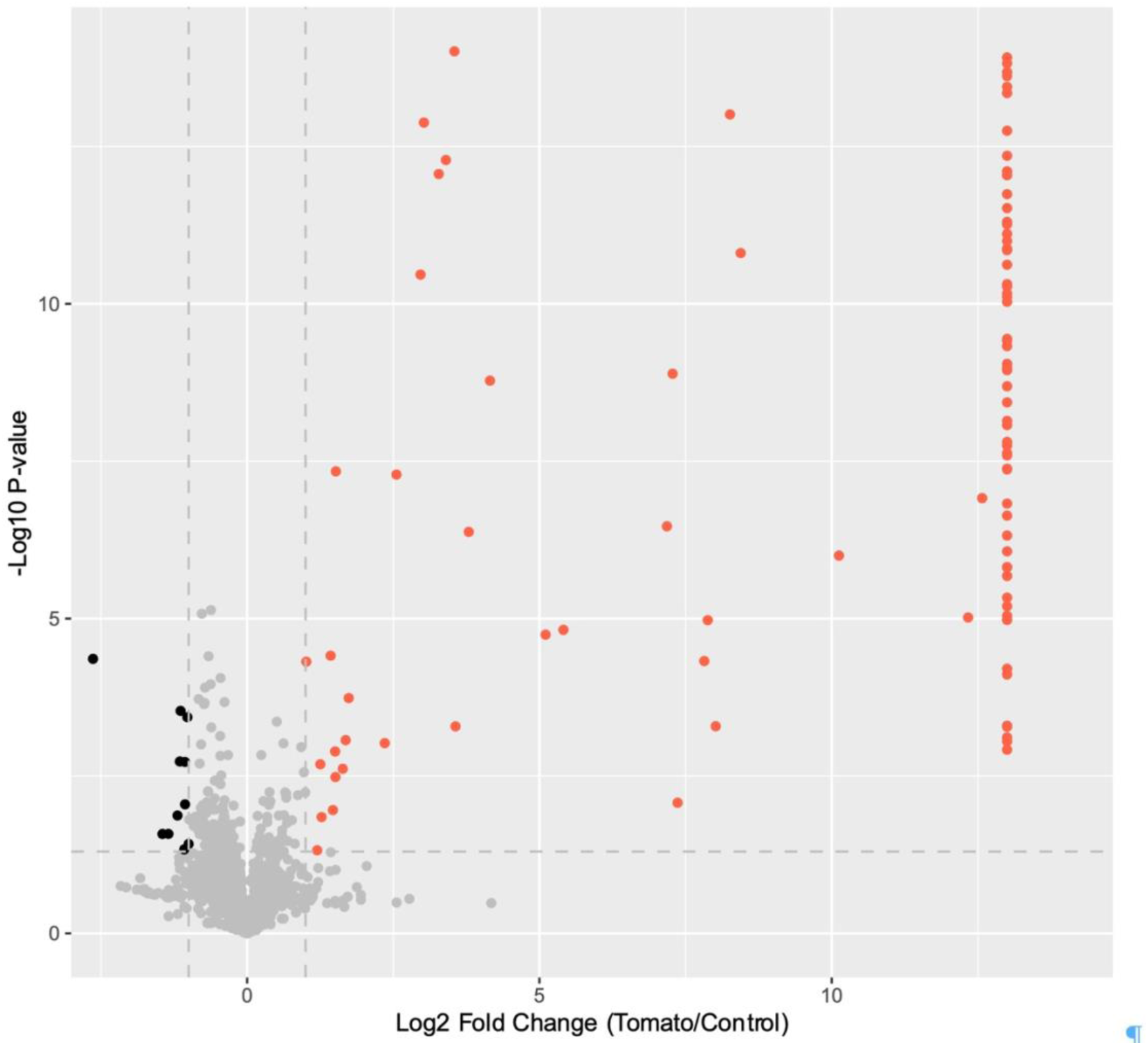
Volcano plot of features detected by untargeted metabolomics (ESI+) comparing mice fed diets enriched with tomatoes to control. Features with a -log10 FDR-adjusted P-value above 2 and a log2 fold change greater than 1 were colored red (higher in mice fed tomatoes). Features with a -log10 FDR-adjusted *P*-value above 2 and a log_2_ fold change less than -1 were colored black (higher in mice fed control diets). Features that were present in tomato and absent from control were assigned a log2 fold change of 13. An interactive version of this plot can be found at www.cooperstonelab.github.io/tomato-liver-omics

A heat map showing significantly different features at an FDR corrected P-value ≤ 0.05 is shown in Figure 3. (Figure 3). The lower portion of the heatmap contains features distinctly absent in control but present in livers of both red or tangerine tomato fed mice. Seventy-five of these features were identified and annotated according to the Metabolomics Standard Initiative guidelines ^[56]^ as tomato steroidal alkaloids and their metabolites (Table 4; Supporting Table 14). Tomatidine and dehydrotomatidine were confirmed by MS/MS spectra and retention time with authentic standards. Seventeen other groups of masses were tentatively or putatively identified as desaturated, hydroxylated, sulfonated, and glucuronidated forms of the steroidal alkaloid aglycones dehydrotomatidine, tomatidine, and esculeogenin B. These data are consistent with our transcriptomics findings where phase I/II genes were differentially expressed from tomato consumption (Tables 2 and 3). A lack of glycosylated steroidal alkaloids, the predominant form in ripe tomato fruit (> 99%) ^[57]^, is consistent with cleavage of sugars prior to uptake of the aglycone into the enterocyte of the small intestine; a documented phenomenon in many other phytochemicals such as polyphenols ^[58]^.

**Figure 3.**
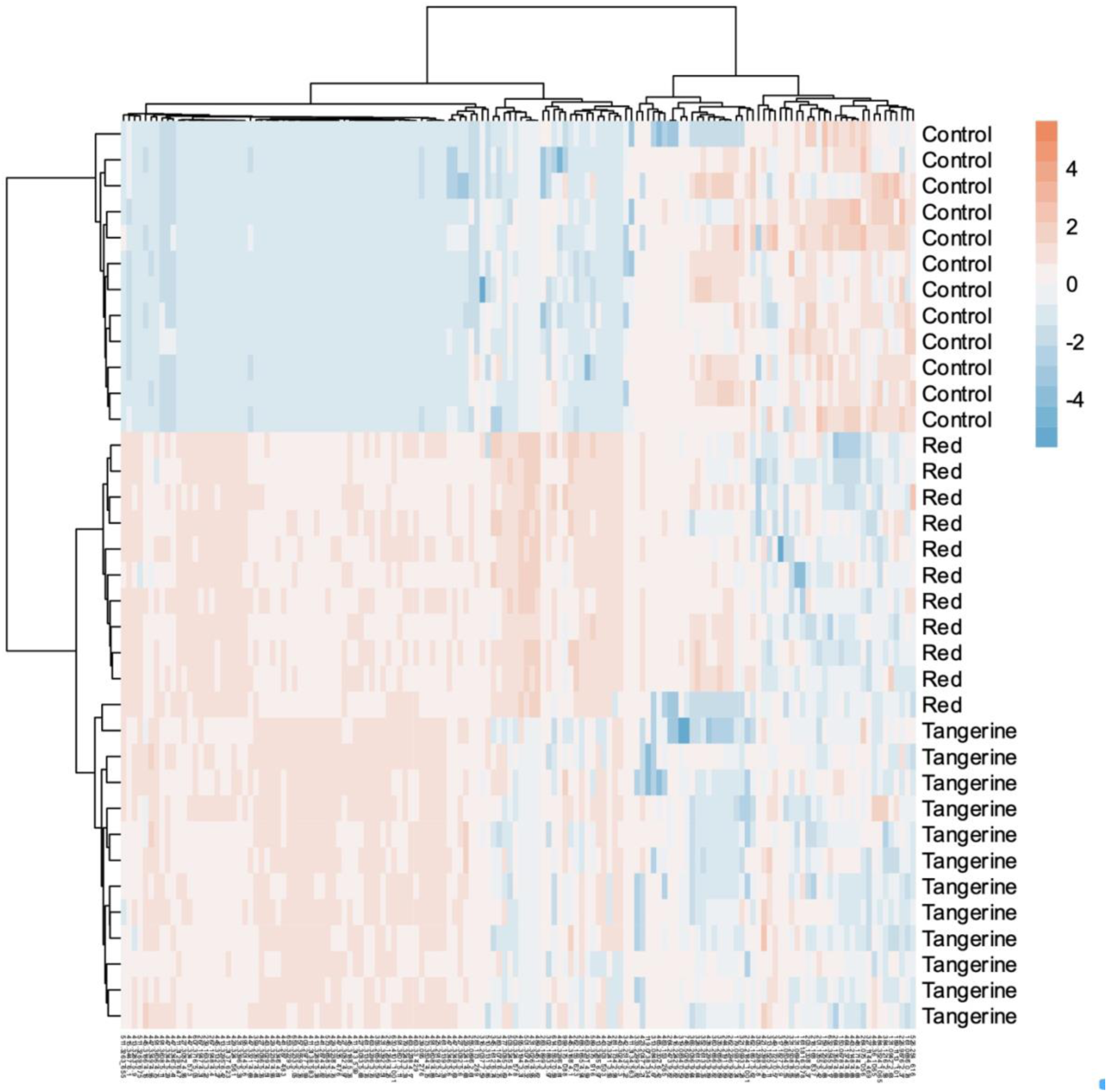
Heatmap of features detected by untargeted metabolomics with ANOVA generated FDR-corrected P-values ≤ 0.05. Hierarchical clustering using Euclidean distances and Ward’s linkage method was used horizontally categorize samples and vertically group features.

**Table 4.** Tomato steroidal alkaloid-derived metabolites present in liver tissue from tomato-fed (red and tangerine) mice.

| Chemical Name | Molecular Formula | Retention Time (min) | Monoisotopic Mass | Observed Mass [M+H] | Mass Error ($\Delta$ ppm)<sup>a</sup> | Distinguishing MS/MS Fragments | ID Level<sup>b</sup> | Additional Isomers<sup>c</sup> |
| --- | --- | --- | --- | --- | --- | --- | --- | --- |
| Didehydro-tomatidine | C<sub>27</sub>H<sub>41</sub>NO<sub>2</sub> | 5.912 | 411.3137 | 412.3216 | -0.24 | 394.3157, 271.2053, 161.1325, 124.1123, 112.0760 | 2 | 4 |
| Dehydro-tomatidine | C<sub>27</sub>H<sub>43</sub>NO<sub>2</sub> | 5.474 | 413.3294 | 414.3366 | 1.45 | 396.2886, 273.2223, 255.2135, 161.1321, 124.1134 | 2 | 4 |
| Tomatidine | C<sub>27</sub>H<sub>45</sub>NO<sub>2</sub> | 6.223 | 415.3450 | 416.3529 | -0.24 | 398.3423, 273.2219, 255.2112, 161.1329, 126.0986 | 1 | 1 |
| Didehydro-hydroxy-tomatidine | C<sub>27</sub>H<sub>41</sub>NO<sub>3</sub> | 5.182 | 427.3086 | 428.3158 | 1.40 | 410.3040, 271.2062, 253.1940, 161.1330, 126.1268 | 2 | 6 |
| Dehydro-hydroxy-tomatine | C<sub>27</sub>H<sub>43</sub>NO<sub>3</sub> | 4.847 | 429.3243 | 430.3318 | 0.70 | 412.3257, 289.2182, 271.2064, 253.0629, 161.1327, 124.1137 | 2 | 7 |
| Hydroxy-tomatidine | C<sub>27</sub>H<sub>45</sub>NO<sub>3</sub> | 5.057 | 431.3399 | 432.3473 | 0.93 | 414.2974, 289.2159, 271.2051, 253.1954, 161.1362 | 2 | 9 |
| Tridehydro-hydroxy-tomatidine | C<sub>27</sub>H<sub>39</sub>NO<sub>4</sub> | 6.604 | 441.2879 | 442.2949 | 1.81 | 424.3126, 289.1929, 267.1182, 253.1974, 161.1321, 159.1151, 157.1020 | 2 | 2 |
| Didehydro-hydroxy-tomatidine | C<sub>27</sub>H<sub>41</sub>NO<sub>4</sub> | 5.3 | 443.3036 | 444.3102 | 0.68 | NA<sup>d</sup> | 3 | 2 |
| Dehydro-dihydroxy-tomatidine | C<sub>27</sub>H<sub>43</sub>NO<sub>4</sub> | 4.258 | 445.3192 | 446.3273 | -0.67 | 428.3205, 289.2156, 253.1958, 161.1332, 157.0996, 128.0707 | 2 | 6 |
| Dihydroxy-tomatidine or<br>Esculeogenin B | C<sub>27</sub>H<sub>45</sub>NO<sub>4</sub> | 4.347 | 447.3348 | 448.3423 | 0.67 | 430.3316, 287.2013,<br>269.1903, 162,1265,<br>126.1284, 124.1092 | 2 | 9 |
| Trihydroxy-<br>dehydro-<br>tomatidine | C<sub>27</sub>H<sub>43</sub>NO<sub>5</sub> | 3.76 | 461.3141 | 462.3213 | 1.30 | NA | 3 | 0 |
| Trihydroxy-<br>tomatidine | C<sub>27</sub>H<sub>45</sub>NO<sub>5</sub> | 4.221 | 463.3298 | 464.3396 | -4.31 | 446.3268, 428.3145,<br>289.2185, 271.2035,<br>253.1955, 156.1025,<br>128.0707 | 2 | 6 |
| Acetoxy-<br>tomatidine | C<sub>29</sub>H<sub>47</sub>NO<sub>4</sub> | 4.75 | 473.3505 | 474.3582 | 0.21 | NA | 3 | 0 |
| Tetra-hydroxy-tomatidine | C<sub>27</sub>H<sub>45</sub>NO<sub>6</sub> | 3.33 | 479.3247 | 480.3321 | 0.00 | NA | 3 | 0 |
| Acetoxy-<br>hydroxy-tomatidine | C<sub>29</sub>H<sub>47</sub>NO<sub>5</sub> | 5.62 | 489.3454 | 490.3529 | 0.00 | NA | 3 | 0 |
| Sulfated<br>hydroxy-tomatidine | C<sub>27</sub>H<sub>45</sub>NO<sub>5</sub>S | 5.52 | 495.3018 | 496.3099 | -0.60 | 416.3528, 298.3426,<br>273.2207, 255.2117,<br>161.1339, 126.1281 | 2 | 0 |
| Glucuronidated<br>dehydro-<br>tomatidine | C<sub>33</sub>H<sub>51</sub>NO<sub>7</sub> | 6.059 | 573.3666 | 574.3736 | 1.39 | 422.6832, 273.2215,<br>255.1923, 244.1200,<br>161.1328, | 2 | 0 |
| Glucuronidated<br>hydroxy-tomatidine | C<sub>33</sub>H<sub>53</sub>NO<sub>9</sub> | 5.139 | 607.3720 | 608.379 | 0.66 | 432.3437, 289.2188,<br>271.2056, 253.1944,<br>161.1318, 126.1279 | 2 | 0 |
| Sulfated<br>Glucuronidated<br>dehydro-<br>tomatidine | C<sub>33</sub>H<sub>51</sub>NO<sub>10</sub>S | 4.972 | 653.3234 | 654.331 | 0.31 | 574.3741, 556.3635,<br>274.1166, 273.2192,<br>269.1901, 255.2149 | 2 | 0 |

<sup>a</sup>Observed mass includes additional proton when calculating mass error
<sup>b</sup>Identification level based on the Metabolomics Standards Initiative <sup>[56]</sup>
<sup>c</sup>Number of additional isomers detected and putatively or tentatively identified for a given chemical feature. Additional retention times can be found in the supporting information.
<sup>d</sup>Not applicable

Common fragments enumerated in Table 4 and Supporting Table 14 match those of authentic standards and previous reports of steroidal alkaloid metabolites ^[16,59–64]^. In many cases, multiple peaks were present for the same *m*/*z*, indicating a series of positional isomers (Table 4; Supporting Table 14). The presence of multiple isomers with unique fragmentation patterns suggests that enzymes involved in xenobiotic metabolism interact with different molecular sites ^[65]^. While the 273 and 255 or 271 and 253 fragments have been previously reported for saturated and desaturated steroidal alkaloids, respectively ^[61,66]^, we also observed masses such as 269 and 251. However, there is not always a perfect relationship between desaturation and the presence of the 271 and 253 fragment as previously demonstrated in hydroxytomatine ^[62]^. The 157 and 124 fragments are hypothesized to be formed from a dehydration reaction of the E and F rings. Example spectra for two representative steroidal alkaloid metabolites are in Figure 4 and positions of double bonds and functional groups were provisionally assigned based on previous reports ^[59,62]^.

**Figure 4.**
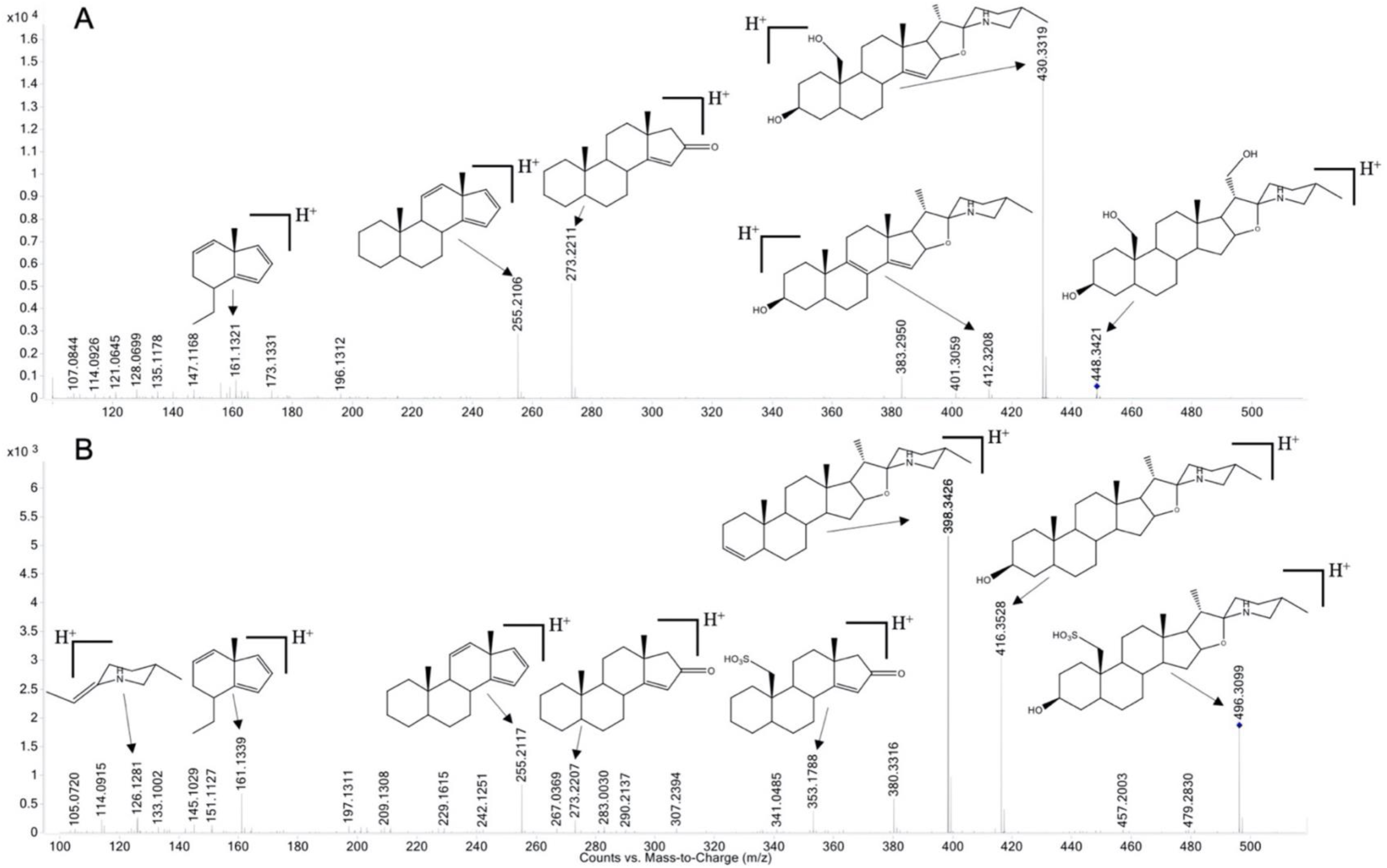
Spectra of dihydroxytomatidine (A) and sulfated hydroxytomatidine (B) generated by MSMS experiments (collision energy: 30 eV). Double bonds and functional groups are provisionally assigned.

Steroidal alkaloids have been previously reported in plasma ^[59]^ and skin ^[16]^ of mice that consumed tomato enriched diets. More recently, metabolites of some tomato steroidal alkaloids have been detected in human urine ^[60,63]^. However, the results here considerably expand this list of identified metabolites. Considering that these chemical features were unique to tomato-fed animals, some may be candidates for tomato consumption biomarkers. A growing body of preclinical evidence suggests that steroidal alkaloids may positively affect health ^[67]^ and their contribution to the bioactivity of tomatoes against chronic diseases in humans warrants further investigation.

## 4 Concluding remarks

Here, we demonstrate that tomato consumption can alter the transcriptional signature of liver tissue and genes related to phase I and II xenobiotic metabolism were most frequently affected. While data generated by RNA-Seq did not clearly discriminate treatment groups, untargeted metabolomics unquestionably separated tomato and non-tomato fed mice based on their chemical profiles; highlighting that significant metabolic differences can exist with subtle transcriptional ones. Through untargeted metabolomics and MS/MS experiments, we confirmed the identity of two steroidal alkaloids and tentatively identified 17 categories of their phase I and II metabolites. These findings greatly expand upon reports of steroidal alkaloids found in mammals where only a few chemical species have been tentatively identified. While once thought to not be absorbed from the diet, we confirmed that steroidal alkaloids are indeed absorbed and metabolized. Given the gene expression signatures we observed and the reported bioactivity of steroidal alkaloids, more research is needed to elucidate the potential impact of tomato steroidal alkaloids on human health and their contributions to the health benefits of tomato consumption.

## Author contributions

M.P.D., D.M.F., S.K.C., and J.L.C. ideated the project, D.M.F. developed and grew tomato material; N.E.M., J.T.A., and S.K.C. provided animal tissues; J.T.A. and M.P.D. extracted RNA and M.P.D. analyzed RNA-Seq data; M.P.D. and J.L.C. carried out untargeted metabolomics, MS/MS experiments, and data analysis; M.L.G. aided in metabolite identification; M.P.D. drafted manuscript; all authors have edited the manuscript; J.L.C. had responsibility for final content.

## Acknowledgements

We thank the Cleveland Clinic Genomics Core (P30CA043703-28) for assistance with library preparation and sequencing, Wilberforce Ouma and Stephen Opiyo for guidance with RNA-Seq analysis, Connor Geraghty for advice regarding gene set enrichment analysis, and J.L. Hartman for help preparing samples for untargeted metabolomics analysis. Financial support was provided by USDA National Needs Fellowship (2014-38420-21844), Hatch funds (OHO01470, OHO01405), USDA-ARS Projects 3092-51000-061-000 (M.P.D) and 3092-51000-061-002S (N.E.M), and Foods for Health, a focus area of the Discovery Themes at The Ohio State University.

## Conflict of interest statement

The authors declare no conflicts of interest.

## Official capacity disclaimer

The findings and conclusions in this publication are those of the authors and should not be construed to represent any official USDA or U.S. Government determination or policy. Mention of trade names or commercial products in this publication is solely for the purpose of providing specific information and does not imply recommendation or endorsement by the U.S. Department of Agriculture. The USDA is an equal opportunity provider and employer.

## Notes

http://www.github.com/CooperstoneLab/tomato-mouse-omics

### Competing Interest Statement

The authors have declared no competing interest.

