## Supplemental Figures for "Transcriptomics and Metabolomics Reveal Tomato Consumption Alters Hepatic Xenobiotic Metabolism and Induces Steroidal Alkaloid Metabolite Accumulation in Mice"

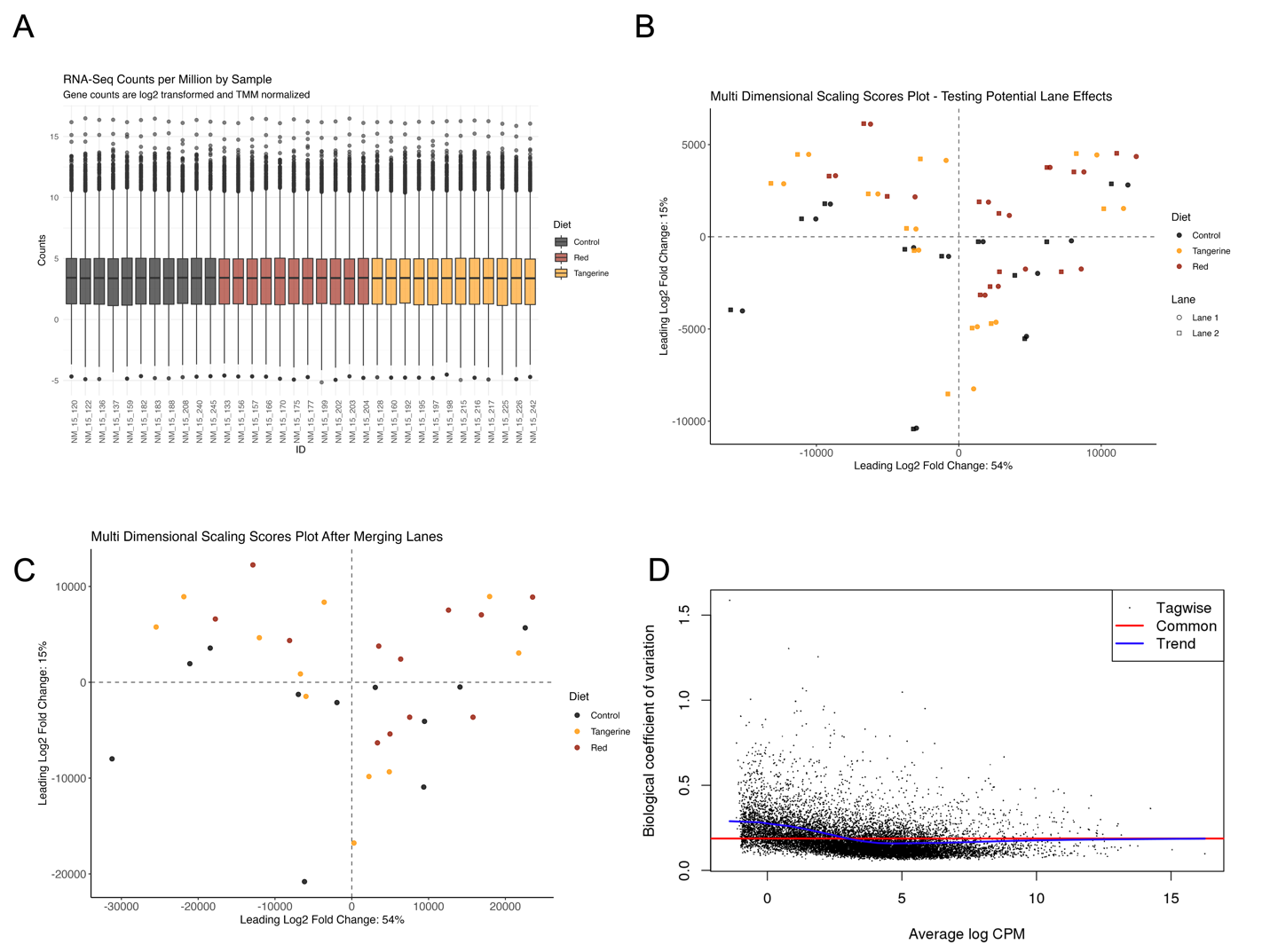
 Figure S1. Boxplots of log2 transformed and TMM normalized RNA-Seq (A); Multidimensional scaling analysis of RNA-Seq data before (B) and after (C) merging data generated on two sequencing lanes; and scatter plot of calculated dispersion of the design matrix used in the analysis of RNA-Seq data (D).

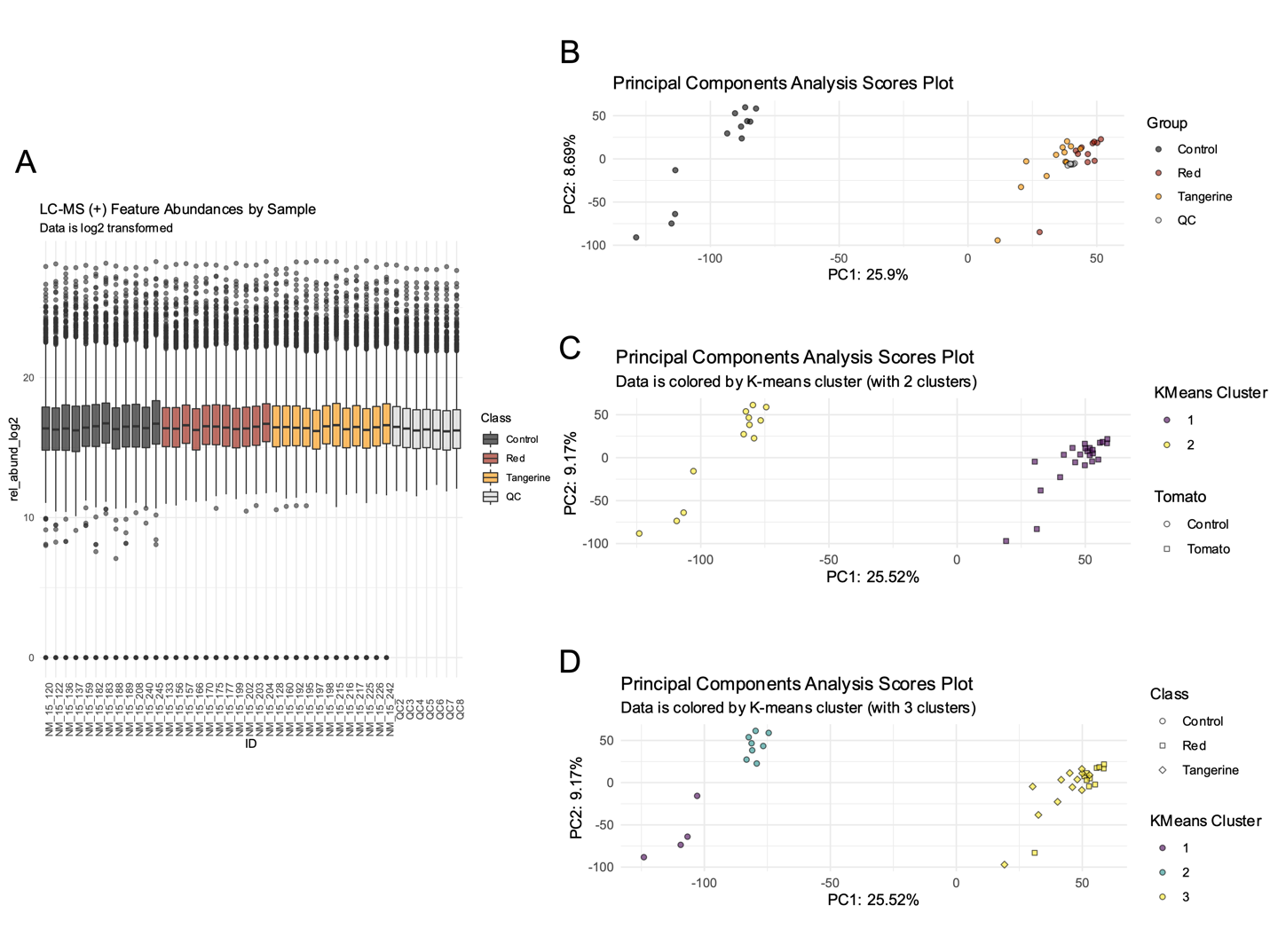

Figure S2. Boxplots of log2 transformed untargeted metabolomics data (A) and principal components analysis scores plot visualizing data structure from untargeted metabolomics (ESI+) data including quality control samples (B). Principal components analysis scores plots with samples colored by assigned k means cluster groups for tomato and control (C) and red, tangerine, and control (D).
